# N-Terminal finger stabilizes the reversible feline drug GC376 in SARS-CoV-2 M^pro^

**DOI:** 10.1101/2021.02.16.431021

**Authors:** Elena Arutyunova, Muhammad Bashir Khan, Conrad Fischer, Jimmy Lu, Tess Lamer, Wayne Vuong, Marco J. van Belkum, Ryan T. McKay, D. Lorne Tyrrell, John C. Vederas, Howard S. Young, M. Joanne Lemieux

## Abstract

The main protease (M^pro^, also known as 3CL protease) of SARS-CoV-2 is a high priority drug target in the development of antivirals to combat COVID-19 infections. A feline coronavirus antiviral drug, GC376, has been shown to be effective in inhibiting the SARS-CoV-2 main protease and live virus growth. As this drug moves into clinical trials, further characterization of GC376 with the main protease of coronaviruses is required to gain insight into the drug’s properties, such as reversibility and broad specificity. Reversibility is an important factor for therapeutic proteolytic inhibitors to prevent toxicity due to off-target effects. Here we demonstrate that GC376 has nanomolar K_i_ values with the M^pro^ from both SARS-CoV-2 and SARS-CoV strains. Restoring enzymatic activity after inhibition by GC376 demonstrates reversible binding with both proteases. In addition, the stability and thermodynamic parameters of both proteases were studied to shed light on physical chemical properties of these viral enzymes, revealing higher stability for SARS-CoV-2 M^pro^. The comparison of a new X-ray crystal structure of M^pro^ from SARS-CoV complexed with GC376 reveals similar molecular mechanism of inhibition compared to SARS-CoV-2 M^pro^, and gives insight into the broad specificity properties of this drug. In both structures, we observe domain swapping of the N-termini in the dimer of the M^pro^, which facilitates coordination of the drug’s P1 position. These results validate that GC376 is a drug with an off-rate suitable for clinical trials.

## Introduction

In late 2019, a respiratory infection initially detected in China, was sparking fear of a viral outbreak [1]. This respiratory infection attributed to severe acute respiratory syndrome coronavirus 2 (SARS-CoV-2), led to an ongoing coronavirus disease 2019 (COVID-19) pandemic with millions infected worldwide (https://coronavirus.jhu.edu/map.html). This respiratory illness was similar to a previous infection by SARS-CoV that led to a SARS outbreak in 2002/3 as well as the Middle East respiratory infection (MERS) outbreak in 2012 [2,3]. All of these outbreaks stem from related betacoronavirus infections, suggesting these strains will likely lead to future viral outbreaks. Vaccines have been developed and will be important for prevention of new infections in the future. However, even with a 95% immunity rate, there will be a significant proportion of people worldwide who will require therapeutic treatment. Antiviral development remains a priority because of importance of immediate mitigation of acute infections, vaccine hesitancy, and the inability to vaccinate some individuals. The outbreak of SARS in 2003 and MERS in 2012 along with the current pandemic reminds us that pan-inhibitors may provide a means for initial control of outbreaks, thereby preventing or quickly controlling pandemics in the future [4].

SARS-CoV-2 is a 30-kb positive-sense single-stranded RNA virus that is translated by the host’s cellular machinery to generate two alternatively spliced long polypeptides, PP1a and PP1ab. These long polypeptides release non-structural proteins (nsps), including the RNA-dependent RNA polymerase, that are essential for viral replication after proteolytic cleavage by proteases from domain nsp3 and nsp5, respectively, a papain-like (PL^pro^) protease and a chymotrypsin-like main protease (M^pro^ or 3CL^pro^)[5]. Similar to SARS-CoV, the SARS-CoV-2 M^pro^ enzyme recognises the sequence of Leu-Gln↓Ser-Ala-Gly, where ↓ marks the cleavage site and this sequence is widely employed for generation of substrates for kinetic analysis and for development of peptidomimetic specific probes and inhibitors [6] [7,8]. The essential role of the M^pro^ in viral replication has resulted in a great deal of crystallographic and *in silico* studies working towards the development of antiviral therapies to treat COVID-19 [9–13].

Proteolytic inhibitors have been used successfully as antiviral therapeutics [14]; for example peptidomimetic inhibitors for the human immunodeficiency virus (HIV) protease and small molecule inhibitors for hepatitis C virus (HCV) protease. The HIV protease inhibitors, along with other drugs, are used in a combination therapy and play a big role in the treatment of symptoms and the subsequent reduction in spread of infection.

It has been recently shown by our group, as well as by other teams, that M^pro^ of SARS-CoV-2 is a promising drug target for the development of SARS-CoV-2 antivirals [10,11,13,15]. We demonstrated that the proteolytic inhibitor GC376 (a bisulphite prodrug) used to treat feline coronavirus infection and its related aldehyde inhibitor, GC373, are effective at decreasing viral load of SARS-CoV-2 in cell culture [13]. These drugs have previously been shown to be effective inhibiting the M^pro^ of picornavirus, norovirus and coronavirus, and furthermore have been validated in animal models for both SARS and MERS [16–18]. Even though we have a considerable understanding of the efficacy of GC376 and GC373 with both SARS-CoV and SARS-CoV-2 M^pro^ [16,17,19–23], detailed mechanistic and functional insight into the inhibitor binding process is still essential for directing broad-spectrum inhibitors in clinical trials. For example, one of desirable features for peptidomimetic proteolytic inhibitors is the reversible nature of binding since it reduces the risk of strong off-target effects and potential toxicity [24,25]. In addition, in light of the new variants, we need a clear understanding of the efficacy of GC373 and GC376 with other coronavirus M^pro^, and importantly a crystal structure of these inhibitors with the SARS-CoV M^pro^ has not been determined.

In this study, we compare inhibition of the M^pro^ of SARS-CoV and SARS-CoV-2 by GC376 using kinetic and structural approaches. We determine K_i_ values are in the low nanomolar range for both SARS-CoV and SARS-CoV-2 M^pro^. After inhibition with GC376, NMR and activity assays demonstrate the reversible nature of inhibition for both proteases. In addition, the restoration of activity of M^pro^ after inhibition reveal a high kinetic and thermodynamic stability for these viral proteases. We determine the crystal structures of SARS-CoV M^pro^ inhibited with the dipeptidyl inhibitor, GC376, and aldehyde form, GC373, both of which reveal a covalent mode of inhibition similar to SARS-CoV-2 M^pro^. We highlight in both structures the role of the N-terminus in stabilizing the S1 subsite from domain swapping, and how this facilitates drug binding. This comparative analysis of M^pro^ from SARS-CoV and SARS-CoV-2 provides additional insight into the mechanism of inhibition by this anti-coronaviral drug.

## Results

### K_i_ values of GC376 inhibition of M^pro^ from both SARS-CoV and SARS-CoV-2 are in nanomolar range

Determining K_i_ values that are reflective of drug binding affinity is a prerequisite for the prediction and evaluation of drug interactions. In our previous report, we determined the half-maximal inhibitor concentrations (IC_50_), values, which describe the functional strength of the inhibitor, the feline drug GC376 with both M^pro^ of SARS-CoV and SARS-CoV-2 [13]. Here we determine K_i_ values for the prodrug GC376 with both M^pro^ of SARS-CoV and SARS-CoV-2. For K_i_ determination, the inhibitory effects of increasing concentrations of GC376 on M^pro^ from both SARS-CoV and SARS-CoV-2 were tested using the synthetic peptide FRET-substrate Abz-SVTLQSG-Y(NO_2_)-R followed by Michaelis-Menten kinetics. Data was plotted as reaction rate versus substrate concentration (primary Lineweaver-Burk plot) and the slopes (Km/Vmax) were determined by linear regression analysis. The slopes were plotted versus the concentration of GC376 to determine the inhibitory constant (K_i_ as y-intercept). The K_i_ for GC376 was 0.02 μM for SARS-CoV M^pro^ and 0.04 μM for SARS-CoV-2 M^pro^, (**Fig 1 and Table 1**).

**Fig 1.**
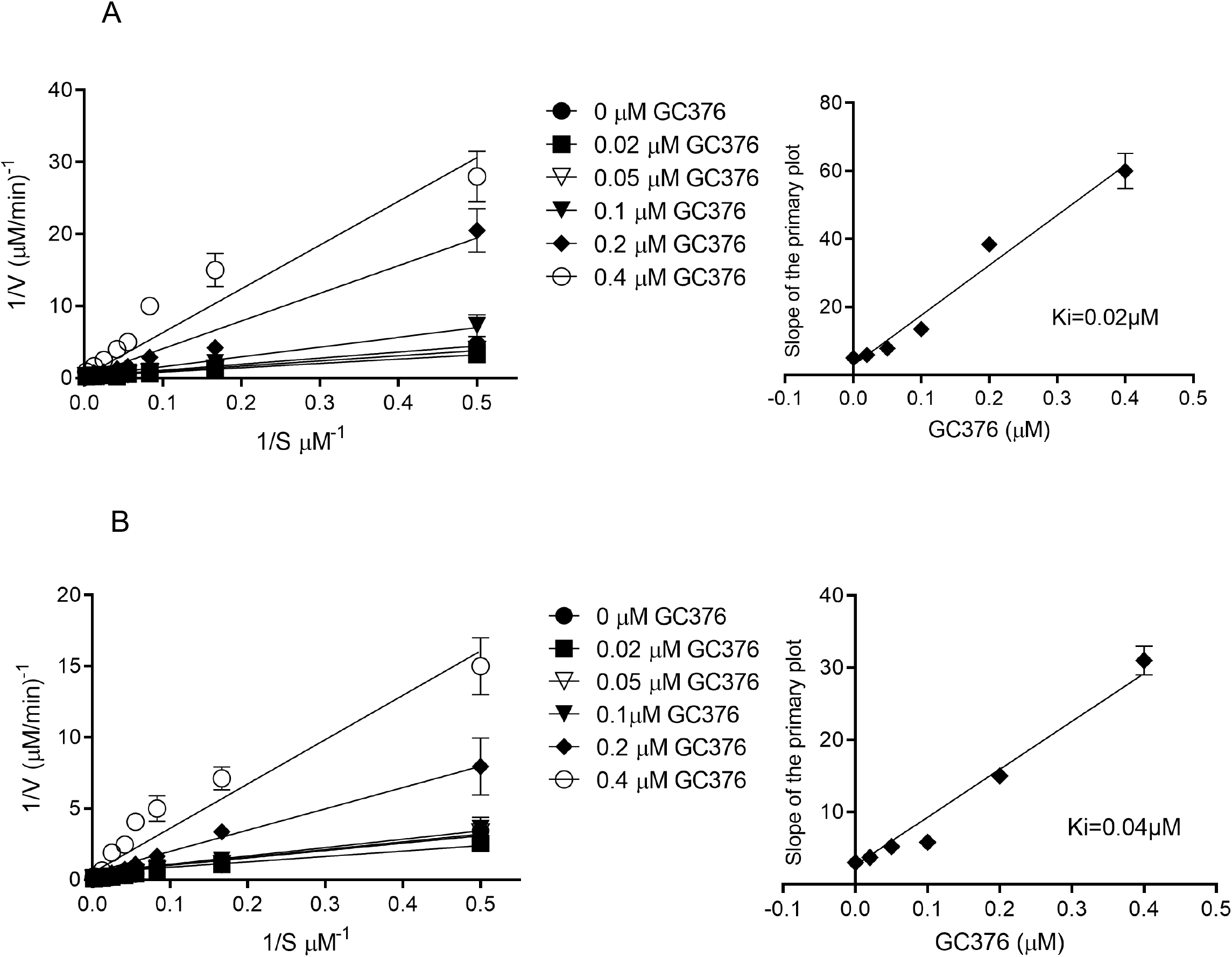
Determination of K_i_ values of GC376 for SARS-CoV M^pro^ and SARS-CoV-2 M^pro^. Lineweaver-Burk plots (left) and the secondary plots of competitive inhibition (right) of SARS-CoV M^pro^ (A) and SARS-CoV-2 M^pro^ (B) by GC376. Data are presented as mean ± SEM, *n* = 3

**Table 1.** Comparison of IC_50_ and K_i_ values between SARS-CoV M^pro^ and SARS-CoV-2 M^pro^ with compound GC376. Data are presented as mean ± SEM, *n* = 3.

| Protease | IC<sub>50</sub>, $\mu\text{M}$ | Calculated K<sub>i</sub>, $\mu\text{M}$ |
| --- | --- | --- |
| SARS-CoV M<sup>pro</sup> | 0.05± 0.01 | 0.02 |
| SARS-CoV-2 M<sup>pro</sup> | 0.19± 0.04 | 0.04 |

### GC376 is a reversible inhibitor with M^pro^ from both SARS-CoV and SARS-CoV-2

An important factor to consider when developing a therapeutic protease inhibitor is the reversibility of compound binding [24]. Irreversible protease drugs can yield long-lasting effects by permanently blocking proteases in cells that are not the intended target and thus causing detrimental consequences resulting in side effects and antigenicity of covalently modified proteins [26]. We previously demonstrated that the bisulfite prodrug GC376 converts to the peptide aldehyde GC373 which interacts covalently with the catalytic cysteine of SARS-CoV-2 M^pro^[13], but did not assess experimentally whether the inhibition was reversible.

Reversibility of GC376 with SARS-CoV-2 M^pro^ was evaluated first by NMR studies using ^13^C-labelled GC373 **(Fig 2A)**. HSQC experiments of samples containing only SARS-CoV-2 M^pro^ **(Fig 2B)**, inhibitor **(Fig 2C)**, or both co-incubated **(Fig 2D)** provided spectra to which the reversibility experiment could be compared. Evidence of binding reversibility was acquired by HSQC experiments conducted on a co-incubated sample containing both enzyme and inhibitor that was subsequently washed with buffer. The subsequent HSQC experiment using this sample showed a disappearance of the NMR signal corresponding to the bound inhibitor (**Fig 2E**). The disappearance of this signal would only be observed in the case of a binding reversion event.

**Figure 2:**
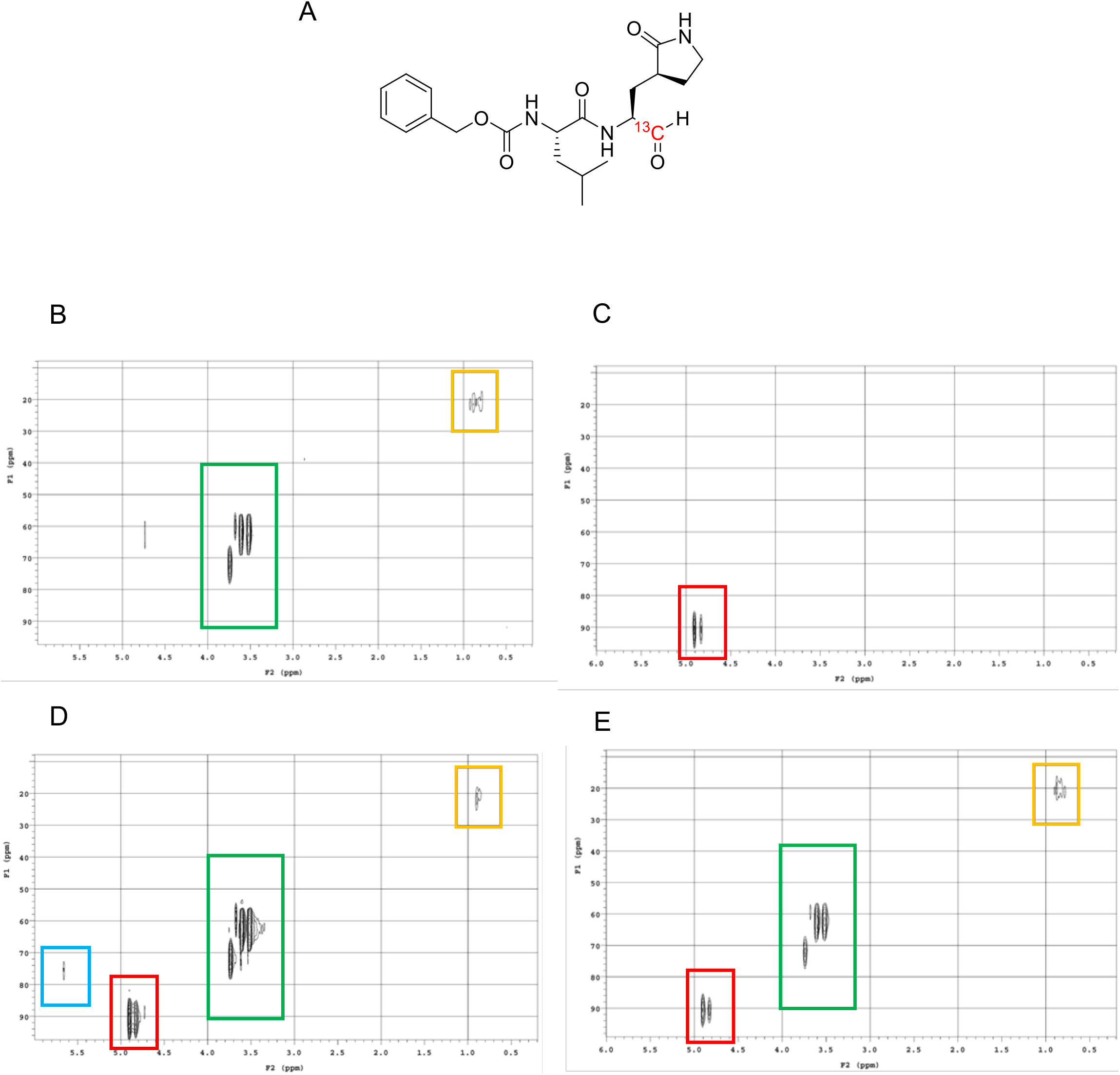
HSQC NMR experiments examining reversibility of GC376 binding. (A) Structure of ^13^C-labelled GC373. (B) HSQC spectra of SARS-CoV-2 M^pro^ in deuterated buffer. (C) HSQC spectra of ^13^C-labelled GC373. (D) Co-incubation of SARS-CoV-2 M^pro^ with ^13^C-labelled GC373. (E) Co-incubated sample after washing step with buffer. Boxes: Blue = bound inhibitor; Red = free inhibitor; Green = DTT (from buffer); Orange = SARS-CoV-2 M^pro^.

We then conducted a detailed study to provide the rate and percentage of reversibility, as well as the comparison of drug behaviour with SARS-CoV M^pro^ and SARS-CoV-2 M^pro^. Reversibility was tested by measuring catalytic activity post dialysis. Incubation of SARS-CoV M^pro^ and SARS-CoV-2 M^pro^ with the GC376 followed by dialysis resulted in increase in enzymatic activity over time, indicative of a reversible dissociation of inhibitor **(Fig. 3)**. We observed a recovery of 10% of activity after 22 hours of dialysis, which reached 30 - 40% of initial activity for SARS-CoV and 40-60% for SARS-CoV-2 after 4 days of dialysis, suggesting over time the substrate competed for the enzyme binding site. To ensure the proteins remained stable over this time period, we also monitored the stability of uninhibited enzymes, which was compared with the activity of recovered enzymes. After 4 days the residual protease activity for the uninhibited M^pro^ of SARS-CoV and SARS-CoV-2 was 30-40%, which allowed us to conclude that the drug was fully reversible.

**Fig 3.**
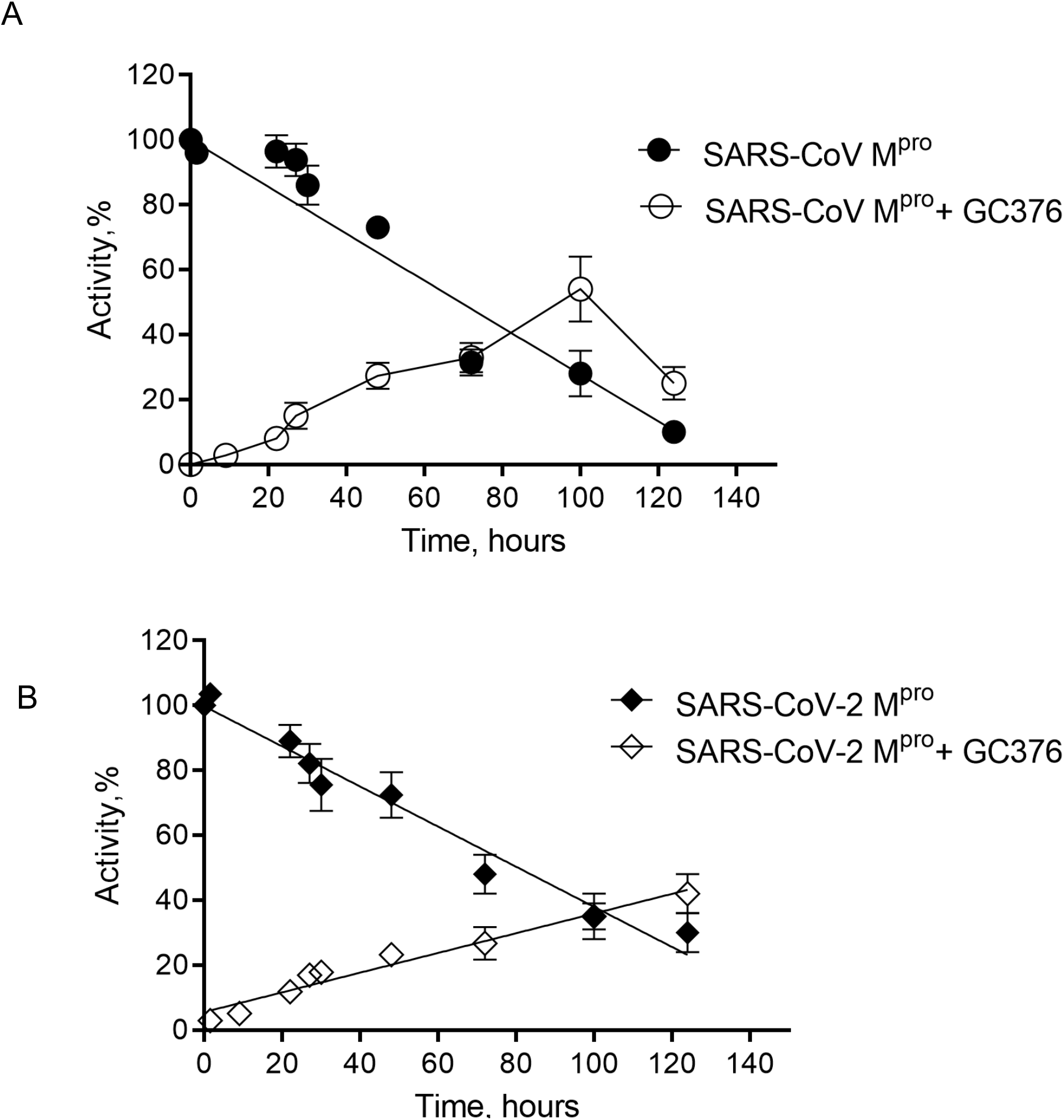
Reversibility of GC376 with SARS-CoV M^pro^ and SARS-CoV-2 M^pro^. The dependence of activity of SARS-CoV M^pro^ (A) and SARS-CoV-2 M^pro^ (B) incubated alone and with the bound GC376 compound on time. Data are presented as mean ± SEM, *n* = 3

### SARS-CoV-2 M^pro^ has enhanced stability compared to SARS-CoV M^pro^

After observing the high kinetic stability of both viral proteases at room temperature, we characterized their thermal stability and assessed their thermodynamic parameters including activation energies of inactivation. Thermal stability is a characteristic used to describe the kinetic stability of enzymes, and many individual proteins or protein complexes are known to have high kinetic stability [27–31]. For viral proteins, particularly the structural ones, this feature is crucial because virus particles must be able to resist harsh environmental conditions until they find a new host to infect and also remain stable during infection [10,13,32]. For example, determination of thermodynamic parameters of the HIV protease in the presence of various inhibitors was used to reveal the differences in protein stability upon forming inhibitor-protein complexes, which informed on inhibitor design [33].

Thermal inactivation of SARS-CoV M^pro^ (**Fig 4A and 4B**) and SARS-CoV-2 M^pro^ (**Fig 4D and 4F**) was studied at the temperature range of 24–70 °C in a time-dependant manner. The semilogarithmic plots of residual activity versus incubation time were linear at all temperatures for both proteins, which was indicative of a simple first-order monophasic kinetic process. From the slopes of semilogarithmic plots inactivation rate constants were calculated and are given in **Table 2**. For both proteases, the rate constant progressively increased with increasing temperatures whereas half-life (*t*_*1/2*_) and the decimal reduction time (Dt), two important parameters used in characterization of enzyme stability, decreased.

**Fig 4.**
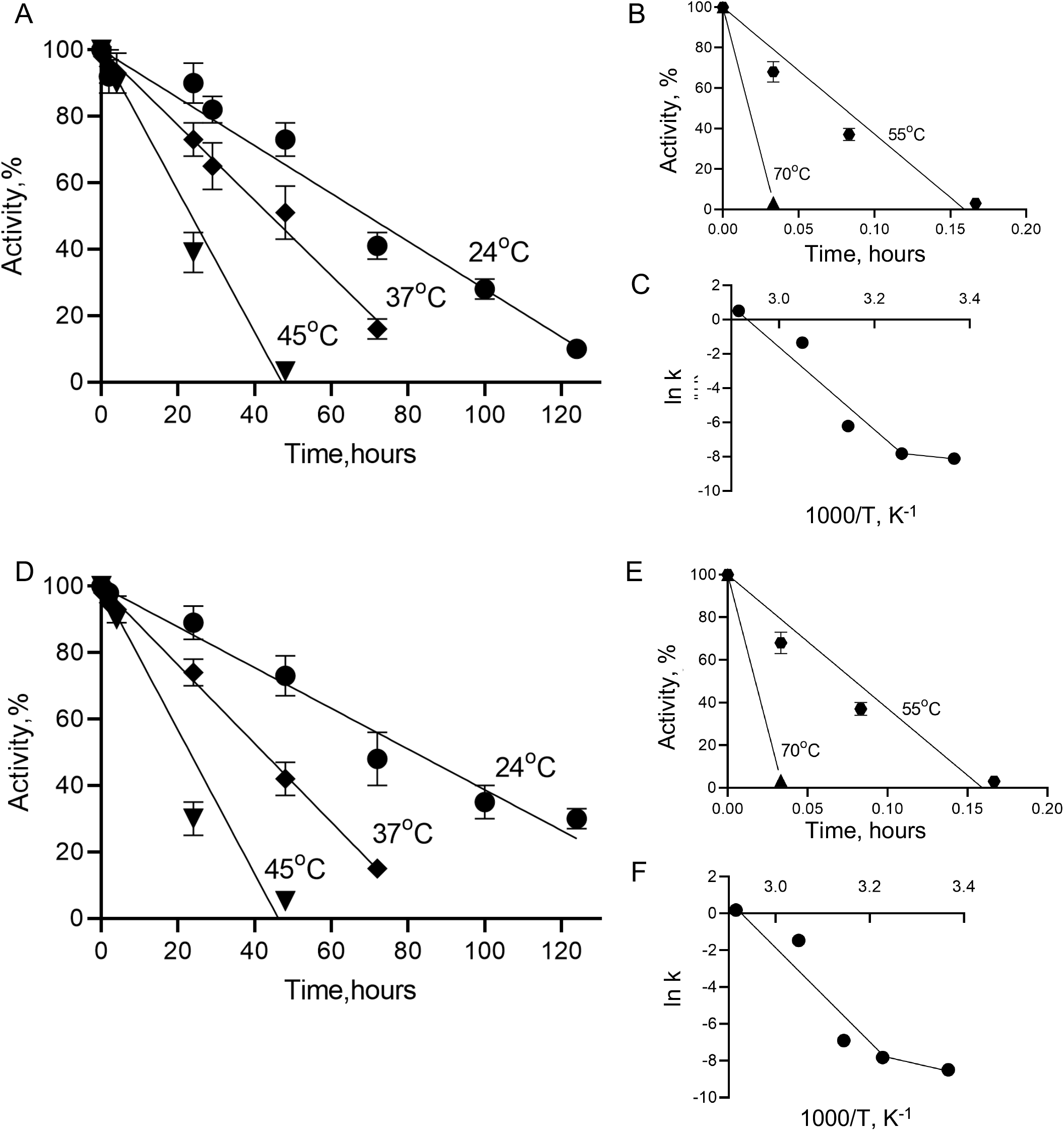
Thermal stability study of SARS-CoV M^pro^ and SARS-CoV-2 M^pro^. Time course of residual activities of SARS-CoV M^pro^ and SARS-CoV-2 M^pro^ in temperature ranges of 24-45°C (A and D, respectively) and in 55-70°C (B and E) and Arrhenius plots for SARS-CoV M^pro^ and SARS-CoV-2 M^pro^ (C and F, respectively). Data are presented as mean ± SEM, *n* = 2

**Table 2.** Thermodynamic parameters for the thermal inactivation of (A) SARS-CoV M^pro^ and (B) SARS-CoV-2 Mpro. *T*, the temperature in oC, *kd*, inactivation rate constant, *t*_*1/2*_, half-life of proteases (i.e., the time after which activity is reduced to one-half of the initial value), *Dt*, decimal reduction time, the time required to reduce the enzymatic activity to 10% of its original value, *ΔG*, activation free energy barrier, *ΔH*, activation enthalpy, *ΔS* activation entropy of thermal denaturation.

| t (°C) | kd (min<sup>-1</sup>) | t<sub>1/2</sub> (min) | t<sub>1/2</sub>(h) | Dt (min) | Dt(h) | $\Delta H$ (kJ/mol) | $\Delta G$ (kJ/mol) | $\Delta S$ (kJ/mol*K) |
| --- | --- | --- | --- | --- | --- | --- | --- | --- |
| 24 | 0.0003 | 2310.5 | 38.5 | 7675.3 | 127.9 | 13.9 | 82.6 | -0.23 |
| 37 | 0.0004 | 1732.9 | 28.8 | 5756.5 | 95.9 | 240.9 | 85.6 | 0.50 |
| 45 | 0.0011 | 630.1 | 10.5 | 2093.3 | 34.9 | 240.9 | 85.2 | 0.48 |
| 55 | 0.318 | 2.2 | 0.04 | 7.2 | 0.121 | 240.8 | 72.5 | 0.52 |
| 70 | 1.67 | 0.4 | 0.007 | 1.4 | 0.023 | 240.6 | 71.2 | 0.52 |

B
| t (°C) | kd (min<sup>-1</sup>) | t<sub>1/2</sub> (min) | t<sub>1/2</sub>(h) | Dt (min) | Dt(h) | $\Delta H$ (kJ/mol) | $\Delta G$ (kJ/mol) | $\Delta S$ (kJ/mol*K) |
| --- | --- | --- | --- | --- | --- | --- | --- | --- |
| 24 | 0.0002 | 3465.7 | 57.7 | 11512.9 | 191.8 | 38.9 | 83.6 | -0.15 |
| 37 | 0.0004 | 1732.8 | 28.8 | 5756.4 | 95.9 | 176.5 | 85.6 | 0.29 |
| 45 | 0.001 | 693.1 | 11.5 | 2302.5 | 38.3 | 176.4 | 85.4 | 0.28 |
| 55 | 0.23 | 3 | 0.05 | 10.01 | 0.2 | 176.4 | 73.4 | 0.31 |
| 70 | 1.2 | 0.6 | 0.009 | 1.9 | 0.03 | 176.2 | 72.2 | 0.32 |

The dependence of inactivation rate constants on temperature was plotted using the Arrhenius equation (**Fig 4C and 4F**), from which apparent activation energies of inactivation (Ea) were calculated. Interestingly, Arrhenius plots for both proteases were not linear and showed upward curvature suggesting two denaturation processes each with its own temperature dependence and activation energy. At temperatures above 37 °C inactivation is a result of protein unfolding with high activation energy, with the rate of this process strongly dependant on temperature. At temperatures of 37 °C and below this rate becomes insignificant and other processes with low activation energy prevail. The activation energies for the high temperature range were found to be high and similar for SARS-CoV M^pro^ (Ea=243.6 kJ/mol) and SARS-CoV-2 M^pro^ (Ea=234.2 kJ/mol). However, for the low temperature range the activation energies were 10—20% of those determined at high temperature, confirming that M^pro^ inactivation involves both high- and low-activation energy processes. Interestingly, the parameters of the inactivation process at low temperature range (24–37 °C) are different for M^pro^ from SARS-CoV and SARS-CoV-2, showing Ea of 16.4 kJ/mol and 41.4 kJ/mol and *t*_*1/2*_ (at 24 °C) of 38.5 h and 57.7 h respectively, suggesting higher stability for SARS-CoV M^pro^.

Determination of all thermodynamic parameters of inactivation can provide further information on enzyme stability. ΔG value, the Gibbs free energy, which is the energy barrier for enzyme inactivation, is directly related to protein stability. We see a significant decrease in ΔG for the temperatures above 55 °C indicating that the destabilization process occurs rapidly in this temperature range (**Table 2**).

To gain a deeper insight into the driving forces of SARS-CoV M^pro^ and SARS-CoV-2 M^pro^ stability, the Gibbs free energy was decomposed into its enthalpic and entropic contributions. Enthalpy, ΔH, measures the number of non-covalent bonds broken during transition state formation for enzyme inactivation, allowing us to compare the energy landscapes of both SARS-CoV M^pro^ and SARS-CoV-2 M^pro^. For temperature ranging from 37 °C to 70 °C we observed consistent high ΔH values, which is in agreement with a temperature-dependent inactivation process. Interestingly, at the 24 °C and 37 °C temperature interval a significant jump in ΔH occured for both proteases, however, with different initial enthalpy values for SARS-CoV M^pro^ and SARS-CoV-2 M^pro^ at 24 °C (13.9 and 38.9 kJ/mol respectively), again highlighting higher stability of latter at physiological temperatures (Table 2). The compactness in the protein molecular structure as well as enzyme and solvent disorder can be inferred through the quantitative analysis of entropy ΔS values [34,35]. Small negative entropy values at 24 °C for both SARS-CoV M^pro^ and SARS-CoV-2 M^pro^ confirmed no disorder in protein structure upon inactivation; however, at higher temperatures all values of ΔS were positive and similar, suggesting that unfolding is a rate-limiting step at this range (**Table 2**).

### Structural comparison of M^pro^ from SARS-CoV and SARS-CoV-2

We previously reported increased catalytic activity of SARS-CoV-2 M^pro^ in comparison to SARS-CoV M^pro^ with the catalytic turnover rate being almost 5 times higher for the former using a FRET-peptide as substrate [13]. We were interested in structural comparison of the M^pro^ from SARS-CoV and SARS-CoV-2, for both apo and drug-bound forms to reveal differences that account for the enhancement in activity. Crystal structures of apo-M^pro^ from SARS-CoV and SARS-CoV-2, and bisulphite prodrug (GC376) and the aldehyde drug (GC373) bound forms were determined. The two proteins share 96% sequence identity with only 12 out of 306 residues being different (**S1 Fig**). Therefore, as expected, there is little change in the overall structures of apo-SARS-CoV and SARS-CoV-2 M^pro^ (**Fig 5**), with an RMSD of 0.6 Å. We observed a new helical feature at ƞ2 (residues 47-50) in SARS-CoV-2, which is unfolded in SARS-CoV, (**S1 and S2 Fig**). It is located at the entrance to the active site, near a non-conserved residue between SARS-CoV, and SARS-CoV-2 (**S2 Fig**). In the GC373-bound form of proteins, however we observed the opposite; this helix is found in the M^pro^ of SARS-CoV but not in SARS-CoV-2 (**S3 Fig**), suggesting a dynamic nature of this structural element.

**Fig 5.**
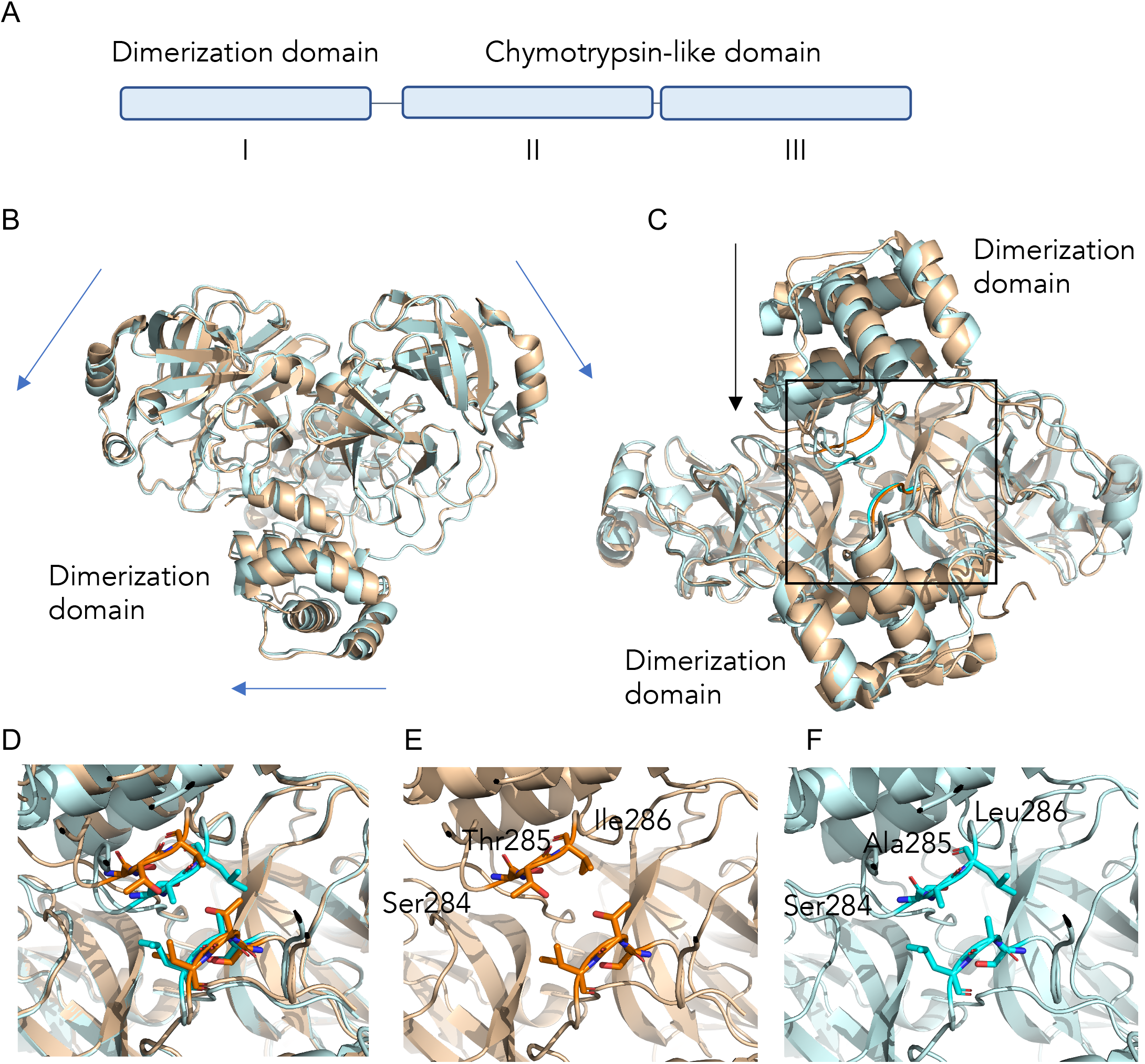
Differences observed between SARS-CoV and SARS-CoV-2 M^pro^ structures. (A) Overall domain organization of M^pro^. (B) Overlay of SARS-CoV M^pro^ (2DUC.pdb) and SARS-CoV-2 M^pro^ (6WTM.pdb) structures reveals small global shifts. (C) The largest structural change is the closer distance between the dimer interface of M^pro^ in SARS-CoV-2 compared to SARS-CoV. (D) A close examination of the dimerization loop in both SARS-CoV M^pro^ (orange) and SARS-CoV-2 M^pro^ (blue). (E) In SARS-CoV M^pro^ a Thr285 in the STI sequence at the dimer interface participates in dimerization via hydrophobic interactions while the M^pro^ in SARS-CoV-2 (F) has an alanine in a SAL motif resulting in a zippered interdigitation of the hydrophobic residues and closer association of the dimerization domains.

Both SARS-CoV and SARS-CoV-2 M^pro^ form dimers, and while monomers have very low activity dimerization is necessary for full enzymatic activity and virulence [36,37]. Comparative analysis of the biological dimer of the two proteases revealed that the main differences are located at the dimer interface. In the M^pro^ of SARS-CoV-2, we observed a slight shift of the chymotrypsin-like domains away from each other, compared to the M^pro^ of SARS-CoV (**Fig 5B**). However, the biggest change is the difference in association between the dimerization domains (**Fig 5C and 5D**). The dimer interface of SARS-CoV and SARS-CoV-2 M^pro^ is facilitated by several interactions between the two protomers, one of which is between the helical domain III of each protomer comprising of residues 284-286, specifically Ser-Thr-Ile (STI) in SARS-CoV M^pro^ and Ser-Ala-Leu (SAL) in SARS-CoV-2 M^pro^. This unstructured loop self-associates between protomers in the dimer. Importantly, this region harbors a non-conservative residue in sequence at the dimer interface, where the Thr285 in SARS-CoV M^pro^ is altered to Ala285 in SARS-CoV-2 M^pro^ (**Fig 5E and 5F**). The SAL-motif forms a tight van der Waals interaction and the residues from each protomer interdigitate to form a complementary interface that readily explains the observed enhanced stability.

### GC376 inhibited forms of SARS-CoV-2 and SARS-CoV M^pro^ reveal a common mechanism of inhibition

We recently presented the structure of GC373 with the SARS-CoV-2 M^pro^ [13]. The structure of SARS-CoV-2 M^pro^ with drug GC373, as well as prodrug GC376 that converts to GC373, reflects the specificity of the enzyme for a glutamine surrogate in the P1 position and a leucine, which is preferred in the P2 position. A benzyl group is in the P3 position. Here we determined the crystal structure of the SARS-CoV M^pro^ with the prodrug GC376 and drug GC373 to examine features that determine its efficacy and compare this with the previously determined SARS-CoV-2 structure (**Fig 6**).

**Figure 6.**
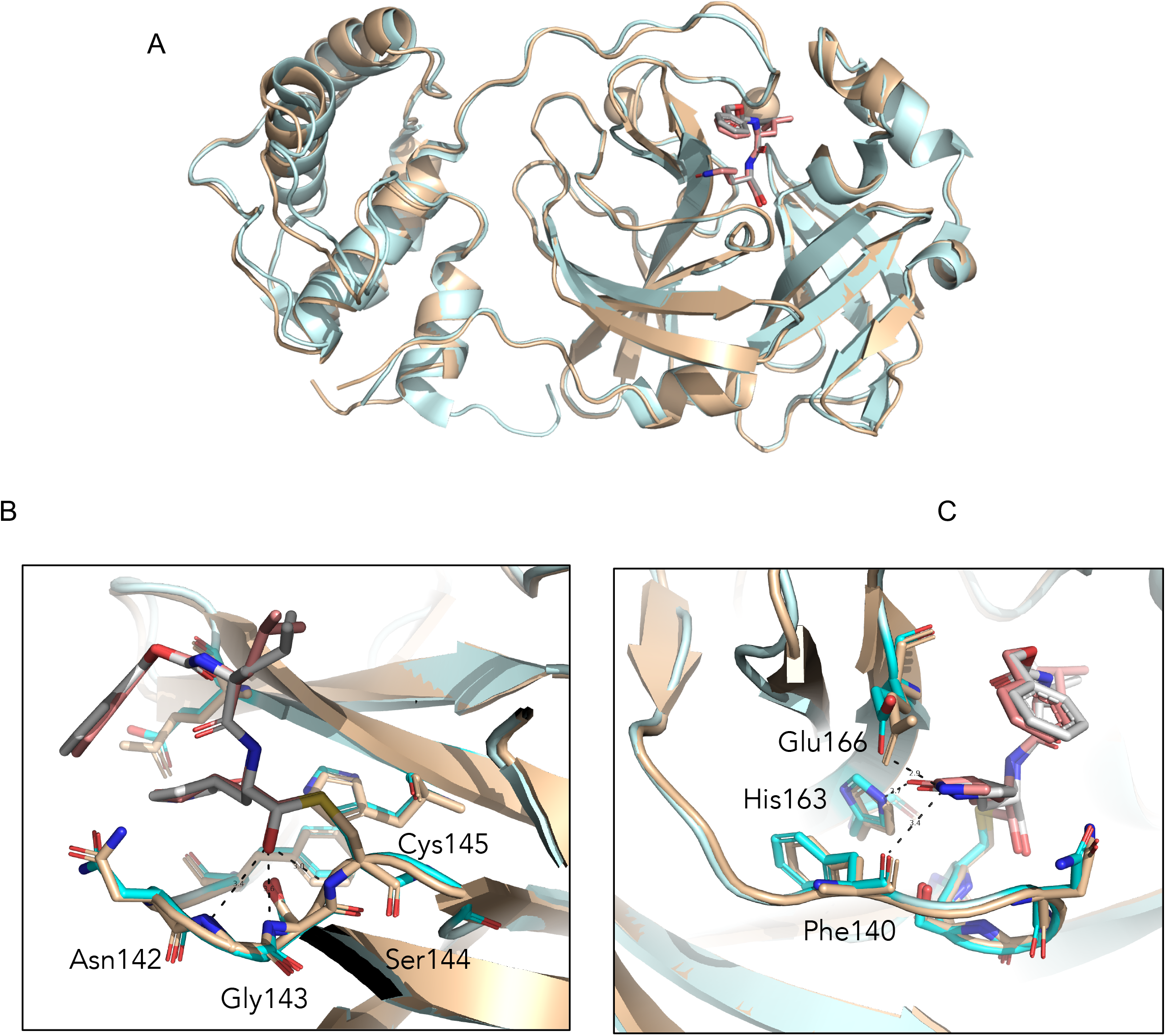
Comparison of SARS-CoV and SARS-CoV-2 M^pro^ structures with GC373 drug. (A). Overall 3-dimensional structures show similarities between SARS-CoV M^pro^ (wheat) and SARS CoV-2 M^pro^ (cyan) with an RMSD of 0.6 Å. GC373 binds covalently with the catalytic Cys145 of the M^pro^ of both SARS-CoV (7LCP.pdb) and SARS-CoV-2 (6TWK.pdb) and shows (B) similar oxyanion hole coordination by Ser144, Gly143 and Asn142 and (C) drug adduct coordination with side chains of His163, Glu166 and backbone of Phe140.

SARS-CoV M^pro^ was incubated with GC373 and GC376, prior to crystallization. The best crystals diffracted to 2.0 Å, and the data was refined with good statistics (**Table 3**). Overall comparison of SARS-CoV M^pro^ and SARS-CoV-2 M^pro^ structures with GC373 showed similar agreements with the apo-M^pro^ structures, with an RMSD of 0.6 Å (**Fig 6**). The drug binding is supported by H-bonding with the main chain of oxyanion hole residues Asn142, Gly143 and Ser144, which are identical for both proteases (**Fig 6B, S4 Fig and S5 Fig)**. A good fit was observed for both the P1 and P2 positions, supported structurally by hydrogen bonding and van der Waals interactions respectively with H-bonds for the P1 position being identical for M^pro^ from SARS-CoV and SARS-CoV-2 (**Fig 6C, S4 Fig and S5 Fig**).

**Table 3.** Data collection and refinement statistics (molecular replacement) for SARS-CoV M^pro^ with drug GC373 and prodrug GC376.

|  | SARS-CoV M<sup>pro</sup> GC373 | SARS-CoV M<sup>pro</sup> GC376 |
| --- | --- | --- |
| PDB entry | 7LCP | 7LCQ |
| **Data collection** |  |  |
| Space group | C2 | C2 |
| Cell dimensions |  |  |
| *a*, *b*, *c* (Å) | 108.28 82.19 54.06 | 108.27, 82.05, 53.82 |
| $\alpha$, $\beta$, $\gamma$ (°) | 90, 104.33, 90 | 90, 104.11, 90 |
| Resolution (Å) | 33.18 - 1.9 (1.968 - 1.9) | 39.19 - 2.15 (2.22 - 2.15) |
| Observations | 236351 (23994) | 110459 (10256) |
| *R*<sub>merge</sub> | 0.053 (1.035) | 0.041 (1.76) |
| *I* / σ*I* | 16.12 (2.08) | 14.7 (0.75) |
| Completeness (%) | 98.04 (97.91) | 96.94 (93.68) |
| Redundancy | 6.6 (6.7) | 4.5 (4.4) |
| CC1/2 | 99.90 (86.70) | 99.90 (60.70) |
| **Refinement** |  |  |
| Resolution (Å) | 33.18-1.90 | 33.19-2.15 |
| No. reflections | 35487 | 24146 |
| *R*<sub>work</sub> / *R*<sub>free</sub> | 19.53/22.08 | 19.58/22.76 |
| No. atoms | 2438 | 2375 |
| Protein | 2358 | 2343 |
| Ligand/ion | 29 | 29 |
| Water | 51 | 3 |
| *B*-factors | 72.13 | 98.79 |
| Protein | 72.31 | 98.98 |
| Ligand/ion | 70.03 | 88.98 |
| Water | 65.22 | 72.66 |
| R.m.s. deviations |  |  |
| Bond lengths (Å) | 0.010 | 0.016 |
| Bond angles (°) | 1.42 | 1.73 |

\*Values in parentheses are for highest-resolution shell. Each data were collected from single crystal

### The N-terminal finger of the M^pro^ stabilizes dimer formation and coordination of the drug GC373

A distinctive feature of M^pro^ dimer is the interaction of N-terminal residues (“N-finger”) of protomer A with residues of domain II of protomer B. In the dimer for both protomers of SARS-CoV-2 M^pro^ and SARS-CoV M^pro^, we observe the N-termini interact with residues near S1 substrate-binding subsite in a hairpin adjacent to the oxyanion hole of the active site (**Fig 7**). The NH-group of Ser1 from protomer A forms strong H-bonds with the carboxylate group of Glu166 (3.1 Å) and the carbonyl of Phe140 (3.3 Å) of protomer B and *vice versa.* This interaction stabilizes the enzyme, assists in the correct orientation of the oxyanion loop and S1 pocket of the substrate binding site, and thus results in enhanced catalytic efficiency, as observed in previous studies demonstrating the native N-terminal serine provides the most efficient enzyme with SARS-CoV M^pro^ [38]. Interestingly, the H-bond distance between the Ser1 (protomer A) and Phe140 (protomer B) is closer in SARS-CoV-2 M^pro^ (3.3 Å) compared to SARS-CoV M^pro^ (5.5 Å) (**Fig. 8**), likely adding to its increased catalytic activity. The proper conformation of S1 pocket is also important for the drug binding and importantly, P1 position of GC373 is also stabilized by hydrogen bonding between the side chain of Glu166 (3.3 Å) and backbone carbonyl of Phe140 (3.3 Å) residues (**Fig 8**). Thus, a hydrogen bond network between the dimer in M^pro^ stabilizes the S1 substrate for substrate binding and hence inhibitor binding.

**Figure 7.**
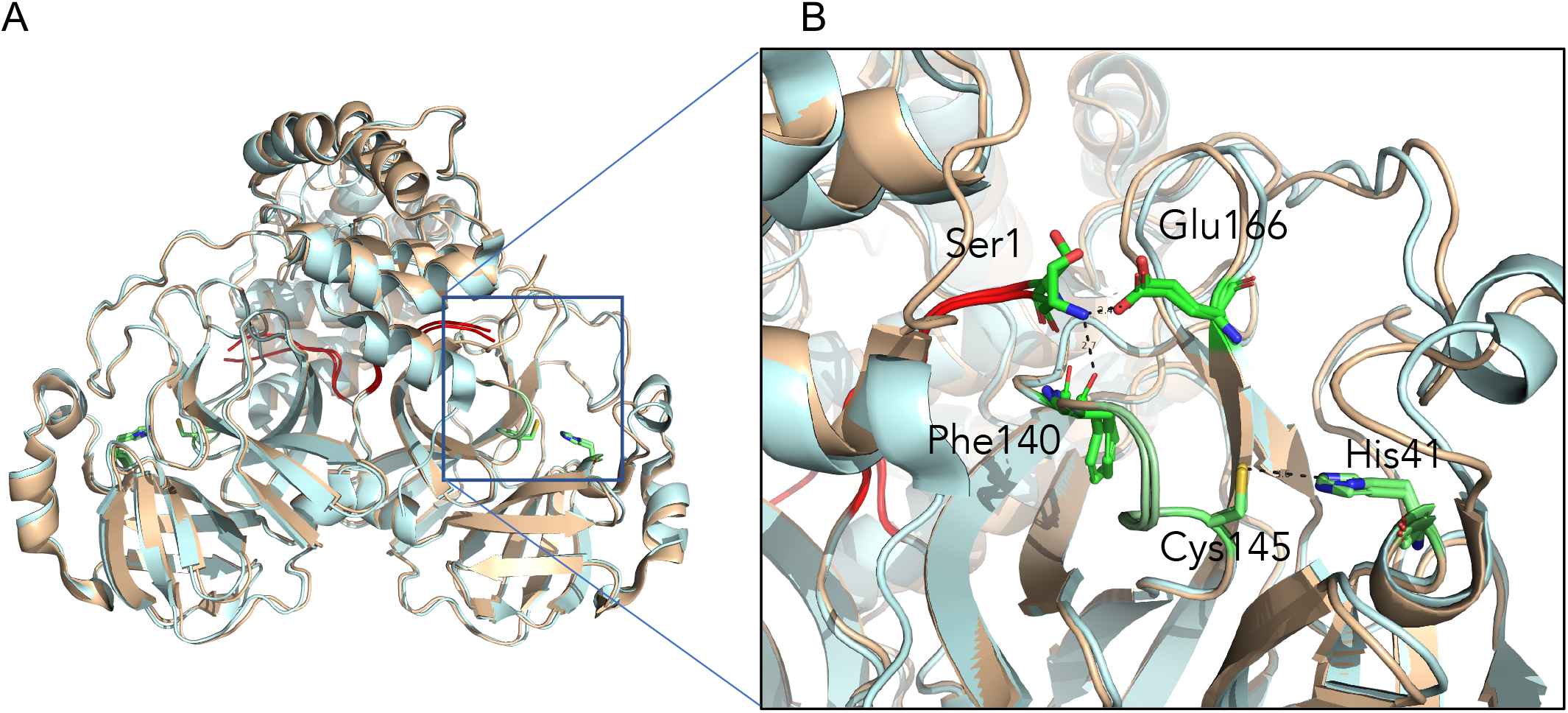
The N-terminus of one protomer interacts with the active site region of the other protomer. (A) Comparison of SARS-CoV M^pro^ (wheat) and SARS-CoV-2 M^pro^ (cyan) structures reveals the N-terminus of each protomer (red) participates in domain swapping in the other protomer. (B) Hydrogen bonding with the N-terminal Ser1 occurs with the side chain of Glu166 and backbone oxygen of Phe140. This influences the region adjacent to the catalytic residues Cys145 and His41, and the oxyanion hole, colored in green.

**Figure 8.**
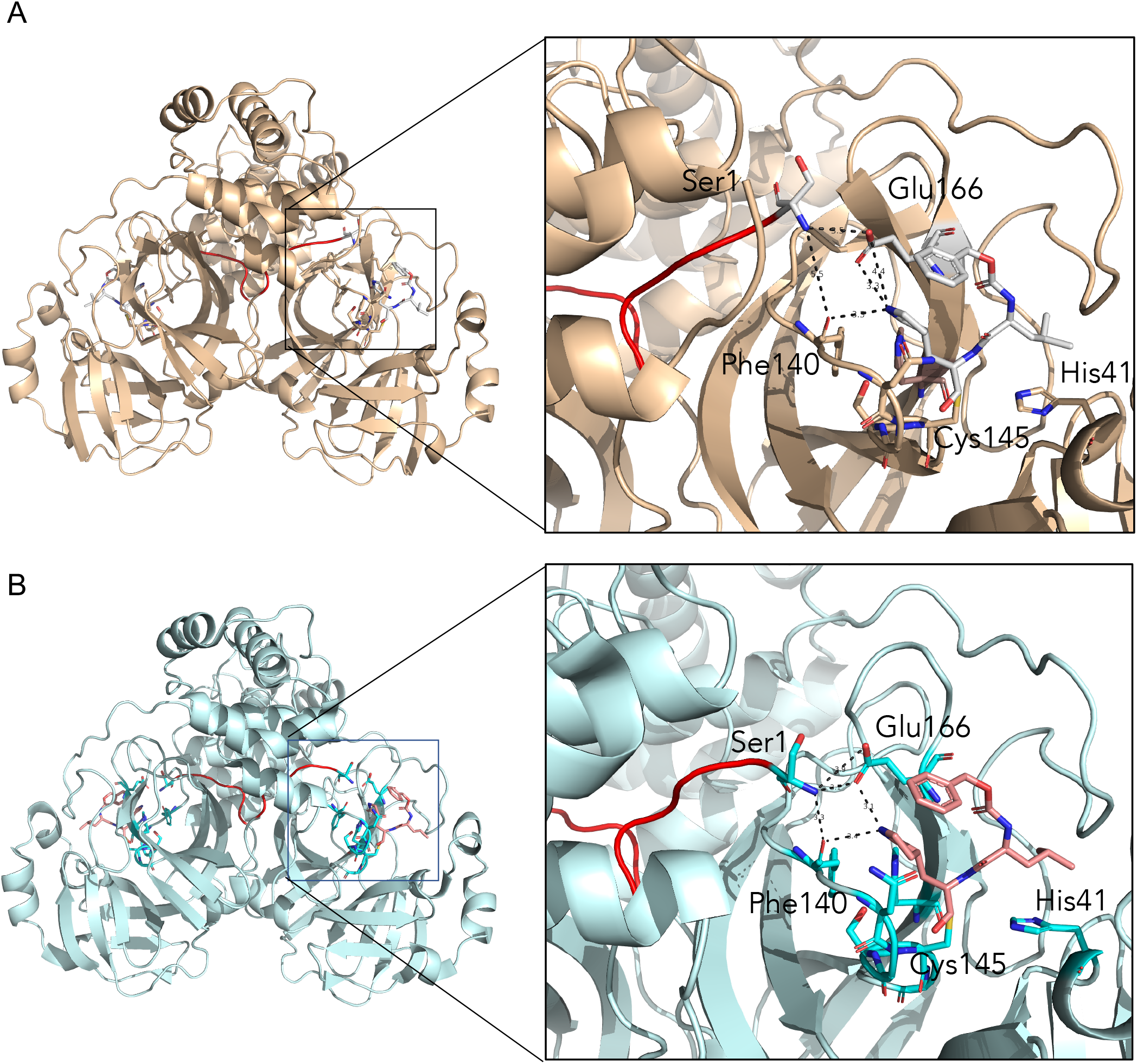
The N-terminal finger of the M^pro^ stabilizes dimer formation and coordination of the drug GC373. The N-finger of M^pro^ facilitates coordination of drug GC373 in both in SARS-CoV (7LCQ.PDB) (A) and SARS-CoV-2 (6WTJ.PDB) (B). Overall of 3-dimensional structures show the N-terminus (red) inserts into the second protomer in SARS-CoV M^pro^ (wheat) and in SARS-CoV-2 M^pro^ (cyan). Both residues that coordinate the N-finger, Phe140 and Glu166 also interact with the P1 position of the drug in M^pro^ of both SARS-CoV and SARS-CoV-2.

Residues adjacent to the N-terminus also play a key role in dimerization, specifically Pro9 and Phe305 from protomer A, which interact with residues Pro122 and Ser123 in a strand on protomer B. We also observe these interactions in all of our SARS-CoV M^pro^ and SARS-CoV-2 M^pro^ structures bound to the inhibitor (**S7 Fig**). Mutation of Pro9 to Thr results in a monomeric species of SARS-CoV-2 M^pro^ [39]. Together this data suggests a strong role for the N-terminus of the protease not only in function and stability, but also with inhibitor coordination.

## Discussion

Here we show that the feline antiviral prodrug GC376 is reversible and inhibits M^pro^ of both SARS-CoV and SARS-CoV-2 with low nanomolar K_i_ values. While IC_50_ values, the concentration of inhibitor at half-maximal inhibition, are very useful during drug development [40], K_i_ values describe precise binding affinity between the inhibitor and enzyme, independent of experimental conditions, and allow for comparisons during structure-activity relationship (SAR) studies. Here we show K_i_ values for GC376 with the SARS-CoV and SARS-CoV-2 M^pro^ to be 20 nM and 40 nM, respectively. These are lower, as expected, when compared to the IC_50_ values of prodrug GC376 (190 nM) and drug GC373 (400 nM) with SARS-CoV-2 M^pro^ [13]. The high degree of sequence identity between the SARS-CoV and SARS-CoV-2 M^pro^ suggests strong conservation in proteolytic inhibition supported by K_i_ values.

K_i_ values for GC376 are in line with K_i_ values of reported proteolytic inhibitors targeting the HCV serine protease and currently being used to treat hepatitis C such as first-generation HCV ns3/4A inhibitors Boceprevir with low nM K_i_ values and second-generation inhibitors with subnanomolar K_i_ values [41]. These drugs are reversible serine protease inhibitors whose development was facilitated by SAR studies [41,42]. Our K_i_ data further supports GC376 being a broad-spectrum inhibitor [16,17,20,22], and demonstrates it is in the inhibitory range to be considered as a viable antiviral for clinical trials.

M^pro^ from SARS-CoV and SARS-CoV-2 have 96% sequence identity and variant residues, with the exception of Ala285 discussed above, are conservative (**S1 Fig**). Therefore, it was not surprising that both proteins revealed similar physical chemical properties such as high thermal stability at temperatures above 37 °C with high activation energies and enthalpy independent of temperature (**Table 2**). However, at physiological temperatures (24-37 °C) we observed a difference in stability between SARS-CoV M^pro^ and SARS-CoV-2 M^pro^, with the latter being more stable, exhibiting higher values of t1/2 (38.5 h for SARS-CoV M^pro^ versus 57.7 h for SARS-CoV-2 M^pro^) and enthalpy (13.9 kJ/mol for SARS-CoV M^pro^ versus 38.9 kJ/mol for SARS-CoV-2 M^pro^). A high ΔH value is usually indicative of a larger number of noncovalent intramolecular bonds, which contribute to protein stability. Therefore, in order to understand what variant residues could be responsible for enhanced stability of SARS-CoV-2 M^pro^ compared to SARS-CoV, we examined the regions with amino acid substitutions more closely.

Both the SARS-CoV and SARS-CoV-2 M^pro^ are dimeric in nature. Early crystal structures of SARS-CoV M^pro^ elucidated how the dimers assemble [7,43] and mutagenesis has revealed that residues at the dimer interface are important for both activity and stability [36,37,44]. From our crystal structures we observe that overall the dimerization motifs of both SARS-CoV and SARS-CoV-2 M^pro^ are very similar, however, one key change at the domain III interface, namely Thr285Ala in SARS-CoV-2 M^pro^, results in a significant alteration in the distance between the domains of the protomers in the SARS-CoV-2 M^pro^ dimer compared to SARS-CoV M^pro^ (**Fig 5**). This mutation leads to residues in the domain III interface forming a hydrophobic zipper clearly aligning the two domains, and thus likely enhancing the t1/2 at low temperatures as we have observed above. The high degree of stability of the enzymes for both SARS-CoV and SARS-CoV-2 is an interesting feature that likely contributes to viral potency.

Another structural feature that might explain the increased activity and stability is a closer association between the N-finger Ser1 and Phe140 in the oxyanion loop in the M^pro^ of SARS-CoV-2 compared to SARS-CoV (**Fig 8**). This interaction plays a critical role for activity since it sustains the correct conformation of the oxyanion loop, therefore precise coordination of the N-finger in both M^pro^ of SARS-CoV and SARS-CoV-2 is a prerequisite for function. Previous work demonstrated that enzymatic activity of SARS-CoV M^pro^ was diminished with non-native affinity tags proving the need for native N- and C-termini [6,38]. The effect was most pronounced with additional residues at the N-terminus, with the activity of the wild-type being 20-fold greater than a variant with an additional glycine at the N-terminus [38].

While GC376 has been crystallized with the main protease of the similar betacoronavirus MERS [18], as well as other viral proteases, including norovirus and porcine diarrhea virus (PEDV) [45], no N-finger association was observed in those crystal structures. This structural motif, however, was observed in a SARS-CoV M^pro^ crystal structure with a Michael acceptor inhibitor, however the N-finger interaction was diminished with the addition of residues at the native N-terminus [38].

We demonstrated that the NH group of Ser 1 donates H-bonds to Phe140 and Glu166, the residues that coordinate the N-termini of each protomer in the dimer. Importantly, these residues also interact with the P1 position of GC373 in both SARS-CoV and SARS-CoV-2, demonstrating a strong hydrogen bond network near the active site, and stabilization of the S1 subsite pocket. This likely contributes to the high K_i_ values for these inhibitors. The precise structural and mechanistic elucidation of the inhibitor-protease interaction and implications for M^pro^ dimerization is paramount for the fine-tuned design of universally active inhibitor drugs. In this regard, the current study provides a rationale for the precise nature of a gamma-lactam group in the P1 position of the GC373/GC376 inhibitor.

With coronavirus outbreaks occurring in 2002, 2015 and 2019, it is clear that broad-spectrum antivirals will be needed for the current pandemic and in the future. The development of antivirals to treat coronavirus infections remains a high priority. By comparing kinetic, thermodynamic, and structural features of M^pro^ from SARS-CoV and SARS-CoV-2 and their binding to GC373/GC376 we revealed distinct supramolecular differences in overall protease properties, yet demonstrate comparable efficacies of GC376 with both proteases. Furthermore, reversible inhibition with the drug further supports the clinical potential of the GC376 compound. The results presented here support the use of GC376 as an antiviral with broad specificity against coronaviruses.

## Methods

### Purification of SARS-CoV M^pro^ and SARS-CoV-2 M^pro^

Purifications of proteases were performed as described earlier [13]. Briefly, pET SUMO (small ubiquitin-like modifier) expression vector (Invitrogen) bearing M^pro^ from SARS-CoV-2 gene with N-terminal His-SUMO tag was transformed into *E. coli* BL21 (DE3), induced with 0.5 mM isopropyl β-d-1-thiogalactopyranoside and the protein was expressed for 4–5 h at 37 °C. After harvesting by centrifugation (4400 × *g* for 10 min at 4 °C) cells were suspended in lysis buffer (20 mM Tris-HCl, pH 7.8, 150 mM NaCl, 5 mM imidazole) and lysed by sonication. The lysate was clarified by centrifugation at 17,000 × *g* for 30 min, and the supernatant was loaded onto Ni-NTA resin column (Qiagen). The resin was washed with 10 column volumes of lysis buffer containing 20 mM imidazole and the fusion protein was eluted with 40–500 mM imidazole in the same buffer. Eluted fractions containing the protein of interest were pooled together and dialyzed against lysis buffer containing 1 mM DTT at 4 °C. The fusion protein was subsequently digested with His-tagged SUMO protease (McLab, South San Francisco, CA) at 4 °C for 1–2 h to remove the SUMO tag and the resulting cleavage mixture was then passed through Ni-NTA resin column. The flow through containing SARS-CoV-2 M^pro^ was collected and further purified using size exclusion chromatography column (G-100, GE Healthcare,) equilibrated with 20 mM Tris, 20 mM NaCl, 1 mM DTT, pH 7.8. Fractions containing the SARS-CoV-2 M^pro^ protein were pooled and concentrated using Amicon Ultra-15 filter with a MWCO of 10 kDa. The plasmid encoding the SARS-CoV M^pro^ with an N-terminal His-tag upstream of a Factor Xa cleavage site was a kind gift of Dr. Michael James. The protein was expressed and purified the same way as SARS-CoV M^pro^-2 but Factor Xa protease (Sigma, Canada) was used (4 °C, overnight) to remove the tag.

### Inhibitor and FRET Substrate Synthesis

Inhibitors GC373 and GC376, and the FRET substrate Abz-SVTLQSG-Y(NO2)-R were synthesized according to methods previously described[13].

### Kinetic experiments

The activity determination of both proteases was performed as previously described[13] using FRET-based cleavage assay with a synthesized fluorescent substrate containing the cleavage site (indicated by the arrow, ↓) of SARS-CoV-2 M^pro^ (Abz-SVTLQ↓SG-Tyr(NO2)-R) in 20 mM Bis-Tris, pH 7.8, 1 mM DTT activity buffer at 37 °C for 10 min. The concentration of proteases was fixed at 80 nM and the range of 0.1–500 μM was used for the substrate. Reactions were started with the enzyme and the fluorescence signal of the Abz-SVTLQ peptide cleavage product was monitored at an emission wavelength of 420 nm with excitation at 320 nm, using an Flx800 fluorescence spectrophotometer (BioTek, USA). The GC376 compound was dissolved in DMSO and used in a concentration range of 0.01–0.4 μM to inhibit both proteases and measure their kinetic parameters. Kinetic data corresponding the interaction of SARS-CoV M^pro^ and SARS CoV-2 M^pro^ with GC376 compound were analyzed using computer-fit calculation (Prism 4.0, GraphPad Software). The slopes of the Lineweaver-Burk plots were plotted versus the inhibitor concentration and the K_i_ values were determined from the x-axis intercept as -K_i_.

### NMR Experiments on Reversibility of Inhibitor Binding

The ^13^C-labelled GC376 inhibitor was synthesized according to previously documented procedures, and initial HSQC NMR experiments involving only enzyme, only inhibitor, and both co-incubated were prepared as previously described [13]. The sample used for the reversibility experiment was prepared by subjecting a previously co-incubated sample containing both enzyme and inhibitor to washing steps with buffer (D_2_O, 50 mM phosphate, pD 7.5 with 20 mM DTT). This involved depositing the sample in an Amicon micro-spinfilter with a 10 kDa cutoff and spinning down the sample at 6600 g for 15 min. The sample was then diluted to 300 μL and the spin down and dilution steps were repeated once more, to a final volume of 300 μL. This sample was then analyzed by NMR in an HSQC experiment, following protocols identical to those previously described [13].

### Reversibility and Stability of 3CL Proteases from SARS-CoV and SARS-CoV-2

Reversibility of 3CL protease inhibition with GC376 was determined by dialysis method. The proteases were incubated with a single concentration (20 μM) of the GC376 compound for 15 min at RT to allow for full inhibition. Then the enzyme-inhibitor mixture was placed in a 6–8 kDa MWCO dialysis membrane (Fisher Scientific, Canada) and dialyzed against 2 L of 50 mM Tris-HCl, pH 7.8, 150 mM NaCl, 5% glycerol, 1mM DTT at RT. The dialysis buffer was changed every 24 hours. Control experiments, which included dialyzing apo-proteases at the same concentration in the same dialysis buffer but different beakers, were performed simultaneously. The aliquots of dialyzing samples were taken out at certain time points and used for activity measurements. The data was represented as a percent of initial protease activity at a zero time point.

The thermal stability was determined by heating 2 μM solution of M^pro^ SARS CoV or M^pro^ SARS-CoV-2 in 50 mM Tris-HCl, pH 7.8, 150 mM NaCl, 5% glycerol, 1mM DTT buffer in a thermostatted water-bath at various temperatures. 30 μl protein samples were taken out at specific time points and immediately incubated on ice until activity measurements were performed as described above. Residual activities were expressed as relative to the maximal activity, which was the activity of proteases at zero time point.

The enzyme inactivation over time is described by a first-order equation:

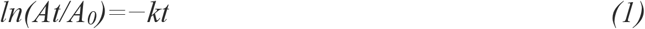

where *A* represents enzyme activity at time *t*, *A*_0_ is the initial activity at time zero, *k* is the rate constant (min^−1^), and *t* is time (min). Inactivation rate constants (*kd*) were obtained from slopes of semi-logarithmical plots of residual activity versus incubation time at each temperature. Calculated rate constants were replotted in Arrhenius plots as natural logarithms of *k* versus the reciprocal of absolute temperature. Arrhenius law describes the temperature dependence of rate constant as

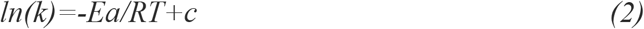

where *Ea* is the activation energy, *R* is the universal gas constant (8.31 J mol^−1^K^−1^), and *T* is the absolute temperature. *Ea* was calculated from the slope of Arrhenius plot.

The half-life of proteases (*t*_*1/2*_), defined as time after which activity is reduced to 50% of initial value [46], was determined as

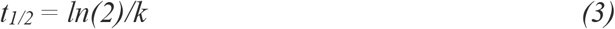

Another common way to present inactivation rate is as *D* value – decimal reduction time, which is the time required to reduce activity to 10% of the original value and calculated as:

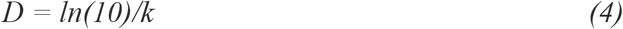

The activation free energy (*ΔG, kJ mol*^−*1*^), enthalpy (*ΔH°, kJ mol*^−*1*^) and entropy (*ΔS°, kJ mol*^−*1*^ *K*^−1^) were determined as

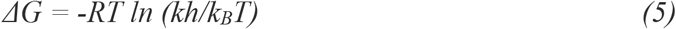

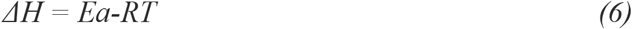

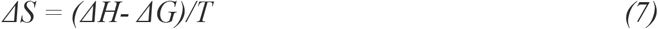

where *h* is the Planck constant (6.626 × 10^−34^ Js) and *k*_*B*_ is the Boltzmann constant (1.38 × 10^−23^ J K^−1^). Experiments were performed in duplicate.

### Crystallization

For crystallization, purified SARS-CoV M^pro^ and SARS-CoV-2 M^pro^ were dialysed against buffer containing 10 mM NaCl and 5mM Tris HCl pH 8.0 overnight at 4 °C. Both proteins were concentrated with a Millipore centrifugal filter (10 kDa MW cut-off) to a concentration of 9 mg/mL. Protein was incubated with 5 molar excess of inhibitor at 4 °C for 2 h prior to crystallization. For SARS-CoV M^pro^, crystals were screened around previously known established conditions [13] with the best crystals forming with vapour diffusion hanging drop trays at room temperature at a ratio of 1:1 with mother liquor containing 10 mM CaCl_2_, 7% PEG 8000, 1 mM MES pH 6.0, 1mM DTT, 3% ethylene glycol and 3% DMSO (Data not shown). For SARS-CoV-2 M^pro^, the protein was subjected to the PACT crystallization screen (Molecular Dimensions, USA), with hits identified in several conditions for both inhibitors. Best crystals were observed with hanging drop trays at room temperature at a ratio of 1:1 with mother liquor 0.2 M Sodium sulfate, 0.1 M Bis-Tris propane pH 6.5, 20% w/v PEG 3350. While the SARS-CoV-2 M^pro^ with ligands crystallize with mother liquid containing 0.2 M Sodium chloride 0.1 M HEPES pH 7.0 20 % w/v PEG 6000. Prior to freezing, crystals were incubated with 15% glycerol as a cryoprotectant for SARS-CoV-2 M^pro^ and 20% ethylene glycol for SARS-CoV M^pro^. Crystals were initially screened at in-house 007 MicroMax (Rigaku Inc) with final data collection at Stanford Synchrotron Radiation Lightsource SSRL, USA, beamline 12-2 with Blu-Ice using the Web-Ice interface [47].

### Diffraction Data Collection, Phase Determination, Model Building, and Refinement

All diffraction data sets were collected using synchrotron radiation of wavelength 0.97946 Å at beamline 12-2 of Stanford Synchrotron Radiation Lightsource (SSRL) California, USA, using a Dectris PILATUS 6M detector. Several data sets were collected from the crystals of SARS-CoV-2 M^pro^ free enzyme as well as with GC376 and GC373 treated. Numerous data sets were also collected for SARS-CoV in the presence of GC376 and GC373. XDS2 [48] and Scala were used for processing the data sets. The diffraction data set of the free SARS-CoV-2 M^pro^ was processed at a resolution of 1.75 Å, in space group P21 (Supplementary Table 1). For the complex of SARS-CoV-2 M^pro^ with GC376 and GC373, the data set collected, was processed at a resolution of 1.9 Å and 2.0 Å and in space group C2 (Supplementary Table 1). All three structures were determined by molecular replacement with the crystal structure of the free enzyme of the SARS-CoV-2 M^pro^ (PDB entry 6Y7M as search model, using the Phaser program from Phenix[49], version v1.18.1-3855). SARS-CoV M^pro^ data were also processed with XDS231 and Scala at a resolution of 2.15 Å and 1.90 Å for GC376 and GC373, respectively, in a space group C2. Ligand Fit from Phenix [50] was employed for the fitting of both inhibitors in the density of pre-calculated map from Phenix refinement, using the ligand code K36. Refinement of all the structures was performed with phenix.refine in Phenix software. Statistics of diffraction, data processing and model refinement are given in (Supplementary Table 1). The model was inspected with Ramachandran plots and all show good stereochemistry. Final models displayed using PyMOL molecular graphics software (Version 2.0 Schrödinger, LLC).

## Competing interests

The authors declare no competing interests.

## Author contributions

J.C.V., W.V. and T.L. contributed to inhibitor synthesis. T.L. contributed to FRET-substrate synthesis. E.A., M.J.v.B., J.L. and C.F. contributed to purified protein. C.F. and E.A. contributed to enzyme kinetics and reversibility studies. M.J.L., H.S.Y. E.A., J.L. and M.B.K. contributed to crystallization and structure determination. W. V. and R. T. M. contributed to labelled NMR studies. M.J.L wrote the initial draft. All authors read and approved the manuscript.

## Acknowledgements

We would like to thank the staff at the Stanford Synchrotron Light Source, in particular Dr. Silvia Russi and Lisa Dunn. M.J.L, J.C.V and D.L.T. acknowledge funding from CIHR and NSERC (COVID-19 SOF-549297-2019). D.L.T acknowledges support from Li Ka Shing Institute of Virology and the GSK Chair in Virology. W.V. was supported by an Alberta Innovates Graduate Scholarship and an Alberta Graduate Excellence Scholarship. T.L. was supported by a CIHR Vanier Scholarship.

Use of the Stanford Synchrotron Radiation Lightsource, SLAC National Accelerator Laboratory, is supported by the U.S. Department of Energy, Office of Science, Office of Basic Energy Sciences under Contract No. DE-AC02-76SF00515. The SSRL Structural Molecular Biology Program is supported by the DOE Office of Biological and Environmental Research, and by the National Institutes of Health, National Institute of General Medical Sciences (P41GM103393). The contents of this publication are solely the responsibility of the authors and do not necessarily represent the official views of NIGMS or NIH.

## Notes

https://www.rcsb.org/structure/unreleased/7LCP

https://www.rcsb.org/structure/unreleased/7LCQ

### Competing Interest Statement

The authors have declared no competing interest.

