## Supplemental figures for "N-Terminal finger stabilizes the reversible feline drug GC376 in SARS-CoV-2 M^pro^"

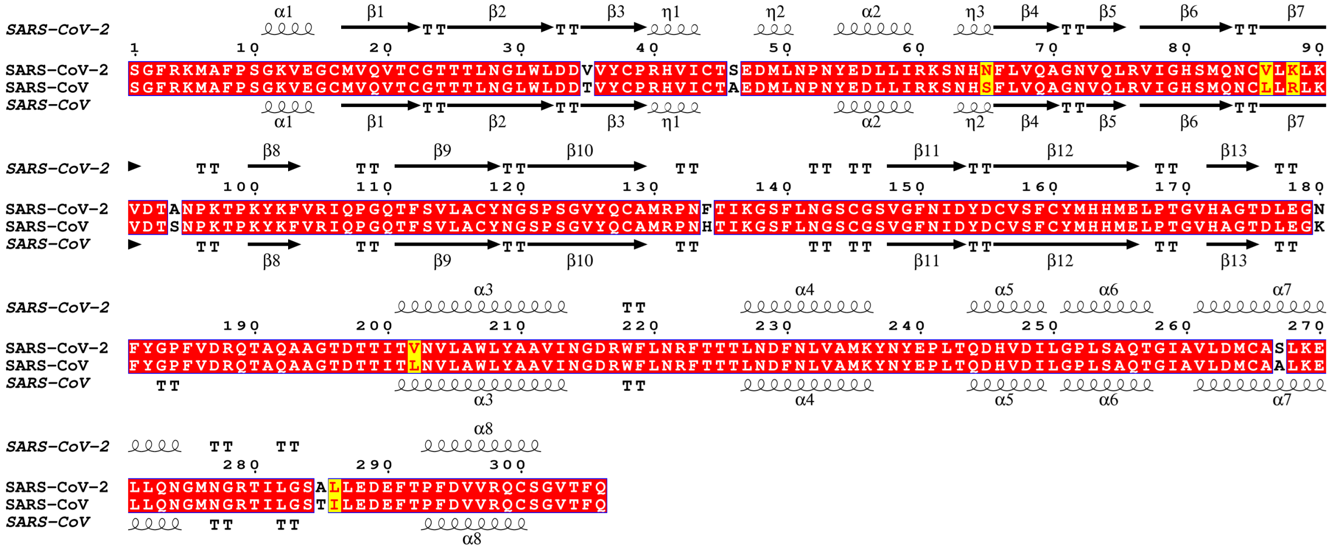


**Supplemental Figure 1**: Secondary structure alignment of SARS-CoV-2 M$^{pro}$ (PDB: 6WTM) and SARS-CoV M$^{pro}$ (PDB: DUC2) using ESPript 3.0.

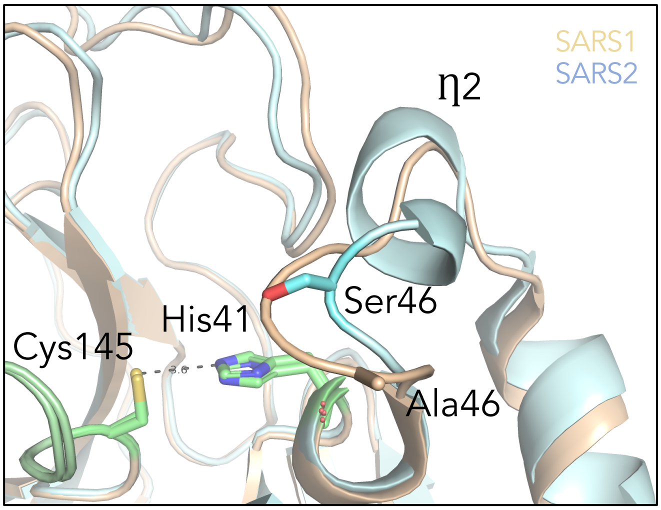


**Supplemental Figure 2. Comparison of SARS-CoV and SARS-CoV-2 M$^{pro}$ structures reveals a new helix.** While overall 3-dimensional structures show similarities between M$^{pro}$ of SARS-CoV (wheat) and SARS-CoV-2 (cyan), a new helix is formed after residue 46, near the Ala to Ser variant. This structural change is adjacent to the active site catalytic dyad, Cys145-His41.

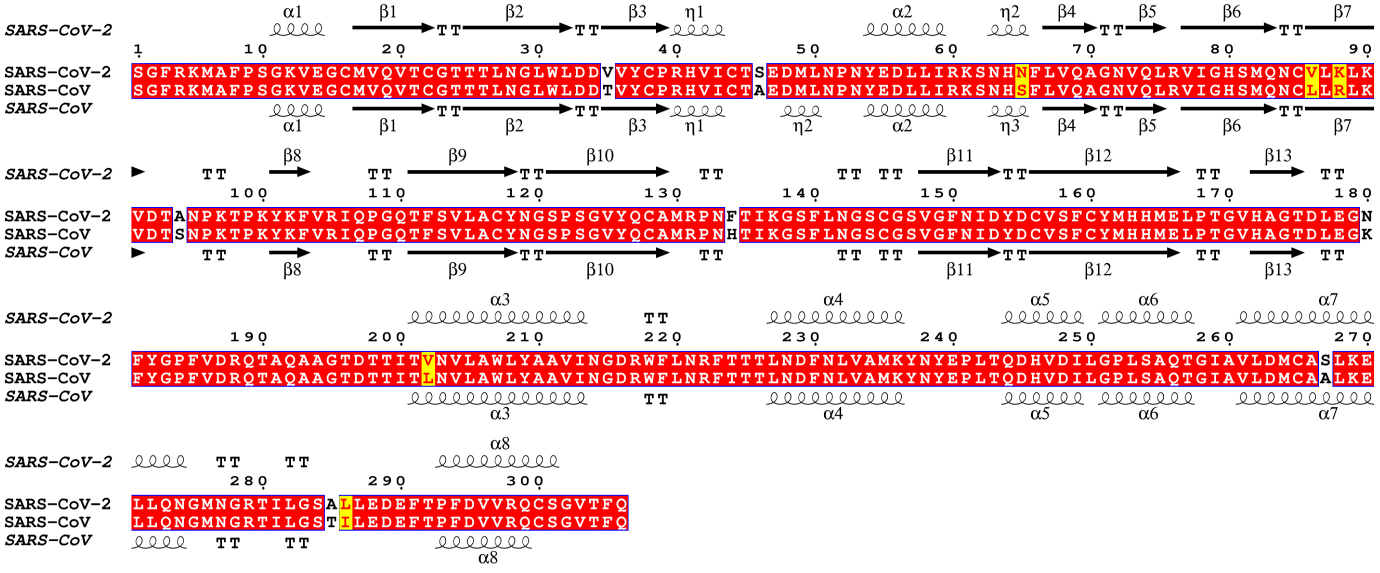

**Supplemental Figure 3:** Secondary structure alignment of SARS-CoV M$^{pro}$ bound to GC373 (7LCP.pdb) and SARS-CoV-2 M$^{pro}$ bound to GC373 (PDB: 6TWK) using ESPript 3.0

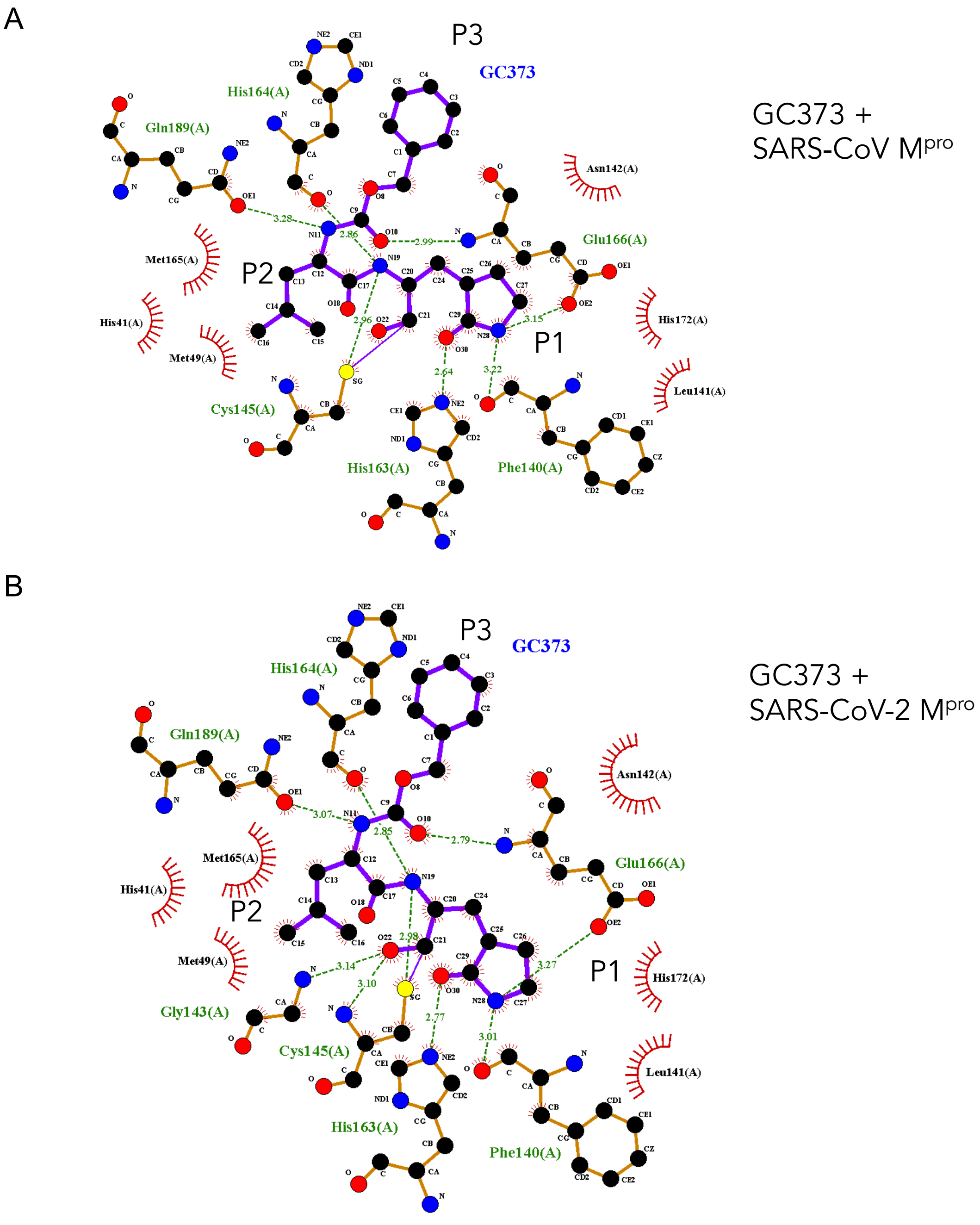


**Supplemental Figure 4**. LigPlot of residues coordinating GC373 to (A) SARS-CoV M$^{pro}$ (PDB:7LCP) and (B) SARS-CoV-2 M$^{pro}$ (PDB: 6WTK).

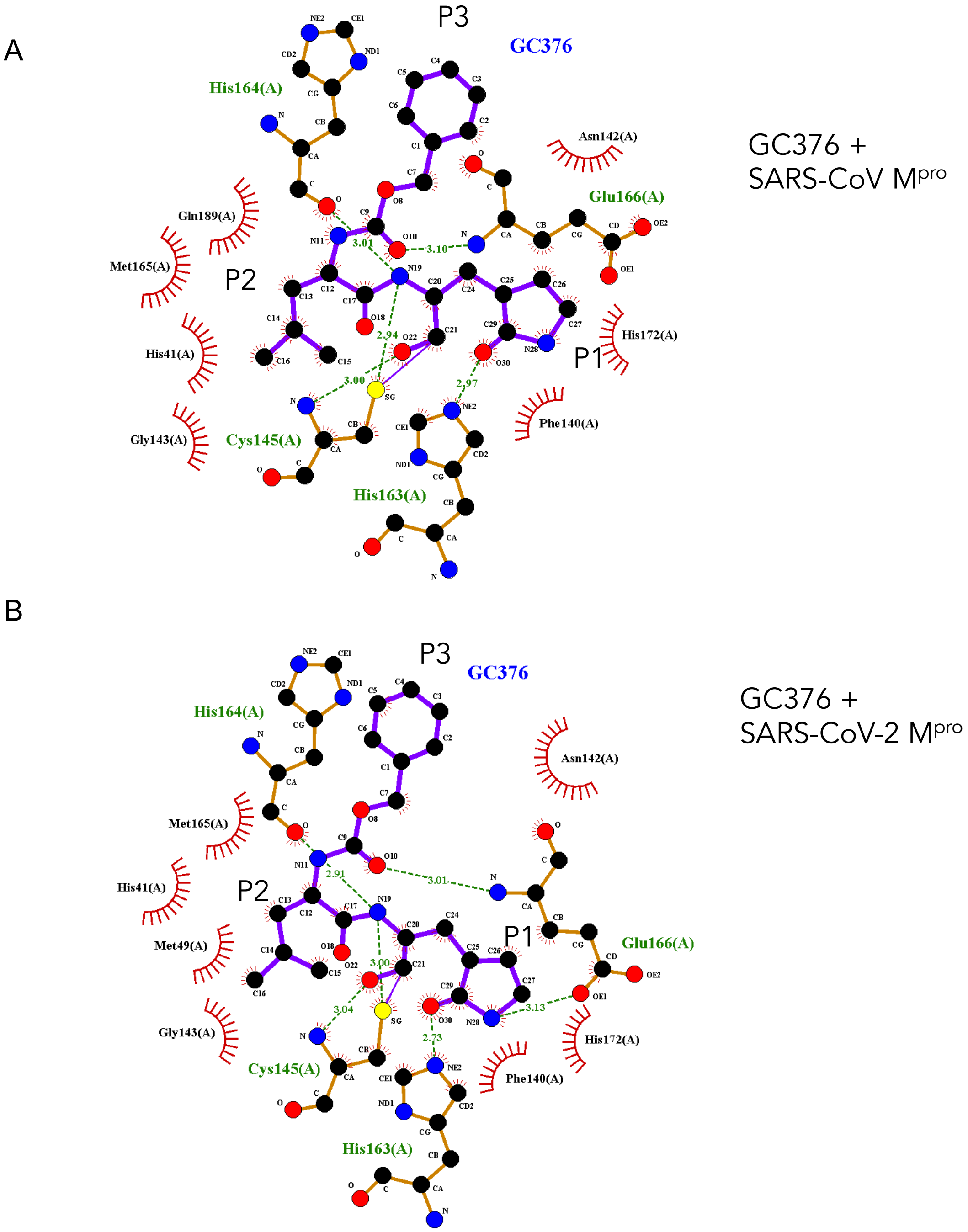


**Supplemental Figure 5.** LigPlot of residues coordinating GC376 to (A) SARS-CoV M$^{pro}$ (PDB: 7LPQ) and (B) SARS-CoV-2 M$^{pro}$ (PDB: 6WTJ).

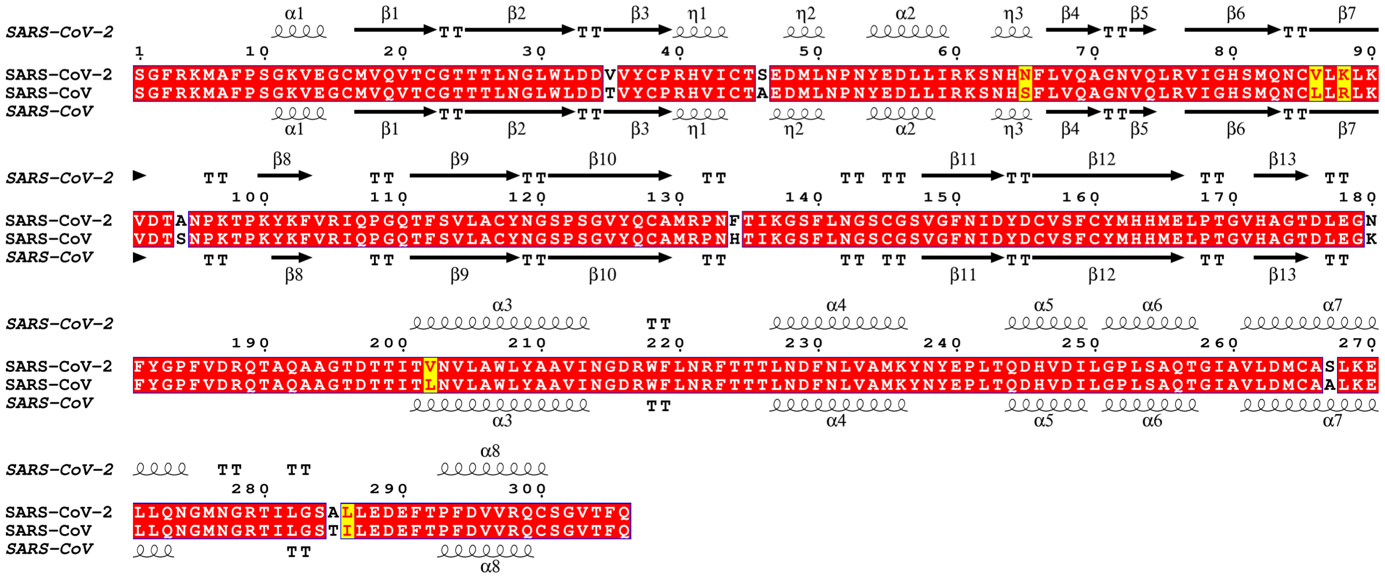

**Supplemental Figure 6:** Secondary structure alignment of SARS-CoV bound to GC376 (7LCQ.PDB) and SARS-CoV-2 bound to GC376 (PDB: 6TWJ) using ESPript 3.0.

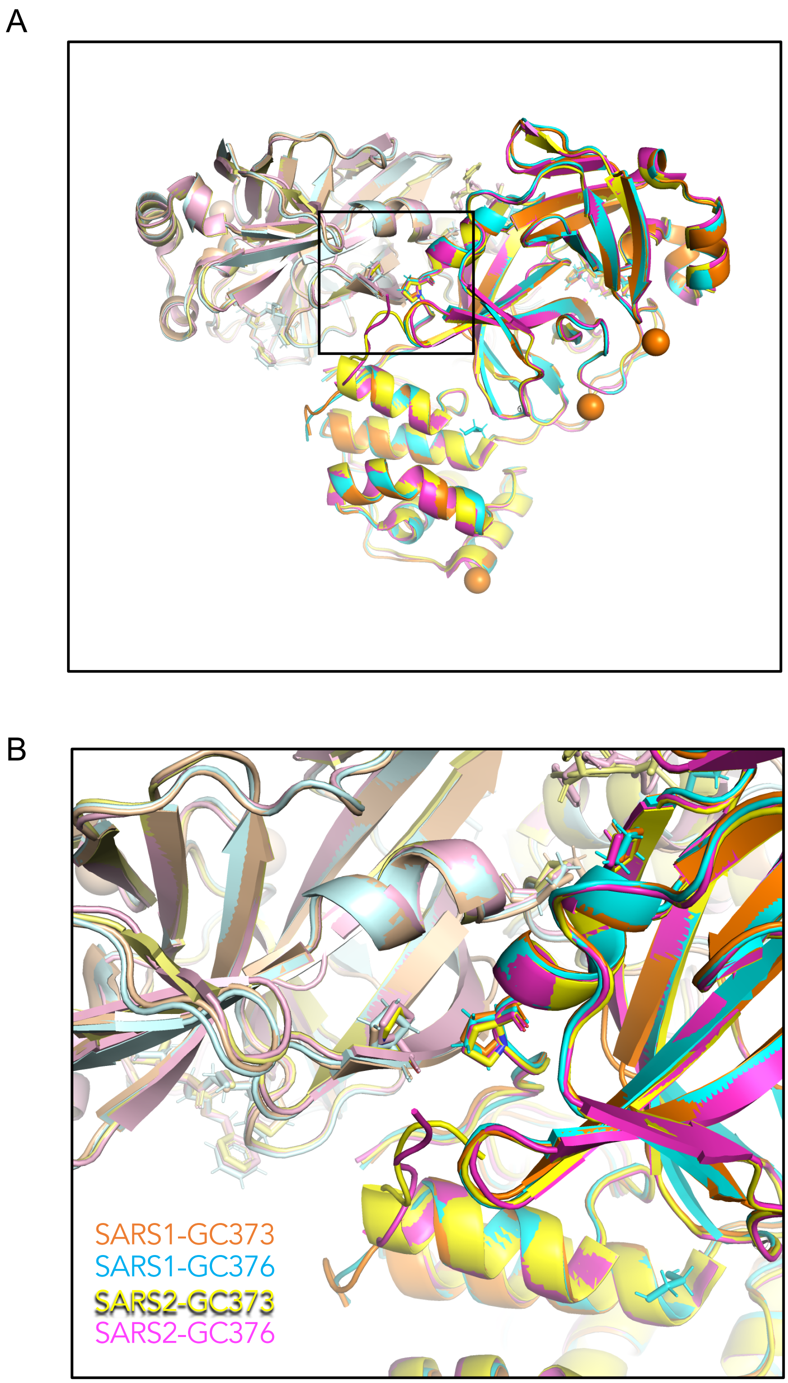


**Supplemental Figure 7. Pro9 (from protomer A) and Pro122 (from protomer B) facilitate dimerization between protomers.** (A) Overall ribbon representation of dimers of GC373 and GC376 bound to SARS-CoV $M^{pro}$ (SARS1) and GC373 and GC376 bound to-SARS-CoV-2 $M^{pro}$ (SARS2) reveals a high structural homology between all structures. (B) A magnified view reveals the Pro9 (from protomer A) and Pro122 (from protomer B) facilitates dimerization.
